# Improved SpCas9 and LbCas12a genome editing systems in *Brassica oleracea* and *Brassica napus*

**DOI:** 10.1101/2022.05.16.492057

**Authors:** Tom Lawrenson, Monika Chhetry, Martha Clarke, Penny Hundleby, Wendy Harwood

## Abstract

We report highly efficient genome editing in Brassica species. We compare the efficiency of targeted mutagenesis using four *Streptococcus pyogenes* Cas9 (*Sp*Cas9) systems in *Brassica oleracea* and *Brassica napus* over 3 target genes and five guides to identify two which show a striking improvement to our first published system (Lawrenson et al., 2015). Targeted mutagenesis occurred in up to 100% of T0 plants with the improved systems, compared to 20% in the original system. This is the only reported comparison of *Sp*Cas9 systems we are aware of in *Brassica* species.

Secondly, we report the first successful use of *Lachnospiraceae bacterium* Cas12a (*Lb*Cas12a) in *Brassica oleracea*. We test three *Lb*Cas12a coding sequences and two guide architectures against one target gene using four guides. From this we identify the best performing combination of our novel, multi-intron coding sequence and a ribozyme flanked guide expression cassette. In this case 68% of T0 plants contained targeted mutations. Heritability of *Lb*Cas12a mutations is shown. We show that our two useful and novel *Lb*Cas12a coding sequences have utility in Brassica species.

## Main text

CRISPR Cas systems are the tools of choice for genome editing in most plant species, with *Streptococcus pyogenes* Cas9 (*Sp*Cas9) and *Lachnospiraceae bacterium* Cas12a (*Lb*Cas12a) the most widely used. We compare our original *Sp*Cas9 system (Lawrenson et al., 2015) to three others and find considerable improvement. We also report the first successful implementation of *Lb*Cas12a in *Brassica oleracea*.

Determination of CRISPR/Cas mutagenesis efficiency was by target loci PCR amplification and Sanger sequencing of amplicons. Plants were scored as positive or negative for mutagenesis.

Our first comparison targeted two unrelated *B. oleracea* genes with *Sp*Cas9 (figure 1a), where guides 1 & 2 (G_1 & G_2) target Bo2g161590 and guides 3 & 4 (G_3 & G_4) target Bo3g005470. System 1 (S1), our original system (Lawrenson et al., 2015), targeted the genes separately with two guides per construct, whilst system 2 (S2) and system 3 (S3) targeted both genes simultaneously with 4 guides per construct. S2 uses an *Sp*Cas9-CYS4 CDS (*At*Cas9Cys4) and a single, multi-guide transcript. S3 uses a plant optimised Cas9 CDS with one intron (PCas9+1int) (Castel et al., 2019) coupled with a tRNA guide architecture (Ma et al., 2019). Figure 1a shows that S2 was not an improvement on S1.

**1a.**
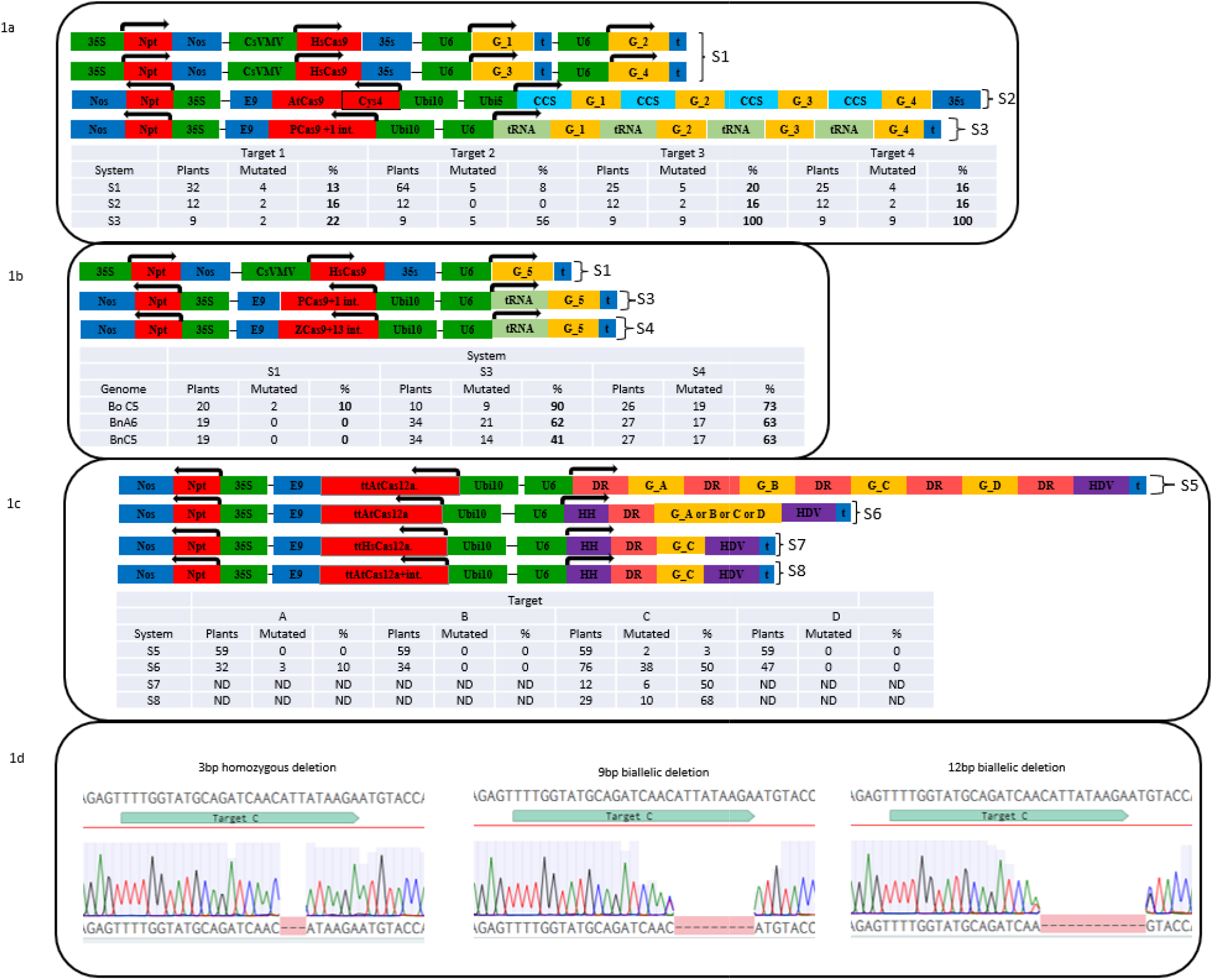
Comparing targeted mutagenesis efficiencies of *Sp*Cas9 systems S1,S2 & S3 on two target genes: Guide 1 and 2 target Bo2g161590, guide 3 & 4 target Bo3g005470. 1b. Comparing targeted mutagenesis efficiencies of Cas9 systems S1,S3 & S4 on *GA4* targets in *B.oleracea* (C5 copy: Bo5g021060) and *B.napus* (A6 copy: BnaA06g10250D; C5 copy: BnaC05G11920D). 1c. Comparing mutagenesis efficiencies of *Lb*Cas12a systems S5,S6,S7 & S8 targeting Bo2g016480. 1d. *Lb*Cas12a mutations inherited in T1. Dark green = promoter; red = CDS; dark blue = terminator; yellow = Guides (G_1-5 sgRNA *Sp*Cas9/G_A-D *Lb*Cas12a spacer); pink = *Lb*Cas12a DR; light blue = Cys4 cleavage site (CCS); Light green = tRNA sequence; purple = self-cleaving ribozyme.

However, S3 did show a striking increase in targeted mutagenesis. Compared to S1, the S3 efficiency was 2 fold, 7 fold, 5 fold and 6 fold greater at targets 1,2,3 & 4 respectively, reaching 100% at targets 3 & 4.

At this point in our investigation a further version of *SpCas9* with 13 introns (ZCas9+13int) was reported (Grützner et al., 2021). This we included as system 4 (S4) in our next comparison, again in conjunction with tRNA guide architecture (figure 1b). We compared S1, S3 and S4 by targeting *GA4* with a single guide (G_ 5) in *B. oleracea* and *B. napus.* In *B. oleracea* guide 5 targets *GA4* on chromosome 5 (Bo5g021060) and in *B. napus* both A6 (BnaA06g10250D) and C5 (BnaC05G11920D) copies are targeted. Previously, 10% of T0 plants were mutagenized at the *B. oleracea* C5 locus using S1 and guide 5 (Lawrenson et al., 2015), but here S1 failed to yield any targeted mutations in *B. napus*. Conversely, S3 was highly effective in both species, with 90%, 62% & 41% of T0 plants carrying targeted mutations respectively at *B. oleracea* C5, *B. napus* A6 and *B. napus* C5. S4 was also highly effective, yielding 73%, 63% & 63% of T0 plants carrying targeted mutations respectively at *B. oleracea* C5, *B. napus* A6 and *B. napus* C5.

High frequency targeted mutagenesis of up to 100% has been reported in *B. oleracea* by Ma et al (Ma et al., 2019) using the tRNA guide architecture and another, intron free version of Cas9 driven by the double 35S viral promoter. It would be interesting to compare this Cas9 to PCas9+1int and ZCas9+13int in *Brassica* species.

Next, we looked at *Lb*Cas12a, and to compare construct designs the gene Bo2g016480 was targeted with 4 guides (A,B,C,D) (figure 1c). System 5 (S5) used an *Arabidopsis* optimised *Lb*Cas12a CDS carrying a ‘temperature tolerant’ D156R mutation (tt*AtCas12a*) (Schindele and Puchta, 2020) coupled to version 1 (V1) guide architecture. Here, 4 guides are driven by one *At*U626 promoter and transcript processing is carried out by Cas12a (Zetsche et al., 2017). A terminal hepatitis delta ribozyme (HDV) prevents spurious guide formation. From 59 S5 T0 plants screened, just two (3%) carried targeted mutations, both of which were located at the guide C target. System 6 (S6) used the same *Lb*Cas12a expression cassette as S5 but has version 2 (V2) guide architecture (Wolter and Puchta, 2019). As S6 constructs each contained V2 single guides, four were made, one each for guides A, B, C & D. Although no mutagenesis was detected at loci targeted by guides B & D, 10% of plants were now successfully mutagenized at locus A and 50% at locus C. By changing the guide architecture alone from V1 to V2 we were able to increase the efficiency of targeted mutagenesis from 0% to 10% at locus A and from 3% to 50% at locus C. We retained V2 guide architecture in system 7 (S7) using guide C in conjunction with a human optimised CDS (Bernabé-Orts et al., 2019). We modified this CDS by inserting the D156R mutation to give tt*Hs*Cas12a. This time 50% of T0 plants carried mutations at locus C indicating that tt*Hs*Cas12a and tt*AtCas12a* work equally well in *B. oleracea.* Taking heed of the improvement made to *Sp*Cas9 by the addition of 13 *Arabidopsis* introns (Grützner et al., 2021) we did the same to tt*AtCas12a*, managing to insert 8 of the same introns whilst retaining a high confidence of splicing to give tt*AtCas12a*+int. When this was used in system 8 (S8), the efficiency of targeted mutagenesis increased to 68% at locus C. The inclusion of 8 introns into tt*AtCas12a* alone increased the efficiency of targeted mutagenesis from 50% to 68%.

To ensure that *Lb*Cas12a derived mutations could be passed to the next generation, two T0 lines with mutations at locus C were analysed in the T1 generation. 24 seeds were germinated for each line and T-DNA free progeny identified using PCR for the NptII marker. From the first line, 9/24 progeny did not contain the T-DNA and all were homozygous for a 3bp deletion at locus C. From the second line 5/24 progeny were T-DNA free, three of which contained 9bp biallelic deletions and two with 12bp biallelic deletions (figure 1d).

In summary we compared four *Sp*Cas9 systems and found two (S3 & S4) worked considerably better than our original S1 system in *B. oleracea* and *B. napus.* We also compared four *Lb*Cas12a systems for the first time in *B. oleracea* and found a human optimised tt*Hs*Cas12a CDS performed equally well compared to an *Arabidopsis* optimised CDS (tt*AtCas12*a). Addition of 8 *Arabidopsis* introns to tt*AtCas12a*, (tt*AtCas12a*+int) resulted in the best performing *Lb*Cas12a CDS. The V2 *Lb*Cas12a guide architecture was superior to V1. We therefore recommend S3 & S4 for *SpCas9* use in *B. oleracea* or *B. napus* and S8 for *Lb*Cas12a use in *B. oleracea.* It is likely that the *Lb*Cas12a system S8 will function in *B. napus.*

## Acknowledgements

Research was supported by the Bill and Melinda Gates Foundation and Foreign, Commonwealth and Development office (ENSA project) and the Biotechnology and Biological Sciences Research Council (BB/P013511/1).

## Author contributions

TL conceived ideas, designed experiments, carried out molecular work and *B. oleracea* transformation. M.Ch. carried out *B. napus* and *B. oleracea* transformation and mutation screening in *B. napus.* M.Cl. and P.H. carried out transformation in *B. oleracea.* T.L. drafted the manuscript. W.H. and P.H. revised the manuscript.

