## Supplemental information for "Improved SpCas9 and LbCas12a genome editing systems in *Brassica oleracea* and *Brassica napus*"

**Guide sequences:**

**G_1: GTCGGAGAAGGGCTGAAGAA**

**G_2: GAAAGAGTTGTAGACTGCGA**

**G_3: GTACGAAGAAGAAACAACCA**

**G_4: GAGTGGAGGGAGAGACAAAA**

**G_5: GTTTTCACTTGCGGCCGGAG**

**G_A: CTTACTAATCTCATTAACAGTTT**

**G_B: CCTCCGCCGGCATTGGAATTCTC**

**G_C: GTATGCAGATCAACATTATAAGA**

**G_D: TGTTCGCGGTATCCCAATCAAGC**

**Kanamycin selection cassette:**

35s promoter

NptII CDS

Nos terminator

gtcaacatggtggagcacgacactctggtctactccaaaaatgtcaaagatacagtctcagaagatcaaagggctattgagacttttcaacaaaggataatttcgggaaacctcctcggattccattgcccagctatctgtcacttcatcgaaaggacagtagaaaaggaaggtggctcctacaaatgccatcattgcgataaaggaaaggctatcattcaagatctctctgccgacagtggtcccaaagatggacccccacccacgaggagcatcgtggaaaaagaagaggttccaaccacgtctacaaagcaagtggattgatgtgataacatggtggagcacgacactctggtctactccaaaaatgtcaaagatacagtctcagaagatcaaagggctattgagacttttcaacaaaggataatttcgggaaacctcctcggattccattgcccagctatctgtcacttcatcgaaaggacagtagaaaaggaaggtggctcctacaaatgccatcattgcgataaaggaaaggctatcattcaagatctctctgccgacagtggtcccaaagatggacccccacccacgaggagcatcgtggaaaaagaagaggttccaaccacgtctacaaagcaagtggattgatgtgacatctccactgacgtaagggatgacgcacaatcccactatccttcgcaagacccttcctctatataaggaagttcatttcatttggagaggacacgctcgagtataagagctcatttttacaacaattaccaacaacaacaaacaacaaacaacattacaattacatttacaattatcgatacaatgaacaagatggattgcacgcaggttctccggccgcttgggtggagaggctattcggctatgactgggcacaacagacaatcggctgctctgatgccgccgtgttccggctgtcagcgcaggggcgcccggttctttttgtcaagaccgacctgtccnggtgccctgaatgaactccaagacgaggcagcgcggctatcgtggctggccacgacgggcgttccttgcgcagctgtgctcgacgttgtcactgaagcgggaagggactggctgctattgggcgaagtgccggggcaggatctcctgtcatctcaccttgctcctgccgagaaagtatccatcatggctgatgcaatgcggcggctgcatacgcttgatccggctacctgcccattcgaccaccaagcgaaacatcgcatcgagcgagcacgtactcggatggaagccggtcttgtcgatcaggatgatctggacgaagagcatcaggggctcgcgccagccngaactgttcgccaggctcaaggcgcggatgcccgacggcgaggatctcgtcgtgacccacggcgatgcctgcttgccgaatatcatggtggaaaatggccgcttttctggattcatcgactgtggccggctgggtgtggcggaccgctatcaggacatagcgttggctacccgtgatattgctgaagagcttggcggcgaatgggctgaccgcttcctcgtgctttacggtatcgccgctcccgattcgcagcgcatcgccttctatcgccttcttgacgagttcttctgagcttgtcaagcagatcgttcaaacatttggcaataaagtttcttaagattgaatcctgttgccggtcttgcgatgattatcatataatttctgttgaattacgttaagcatgtaataattaacatgtaatgcatgacgttatttatgagatgggtttttatgattagagtcccgcaattatacatttaatacgcgatagaaaacaaaatatagcgcgcaaactaggataaattatcgcgcgcggtgtcatctatgttactagatcga

**S1 *Hs*Cas9 cassette:**

CsVMV promoter

*Hs*Cas9 CDS

35S terminator

ccagaaggtaattatccaagatgtagcatcaagaatccaatgtttacgggaaaaactatggaagtattatgtaagctcagcaagaagcagatcaatatgcggcacatatgcaacctatgttcaaaaatgaagaatgtacagatacaagatcctatactgccagaatacgaagaagaatacgtagaaattgaaaaagaagaaccaggcgaagaaaagaatcttgatgacgtaagcactgacgacaacaatgaaaagaagaagataaggtcggtgattgtgaaagagacatagaggacacatgtaaggtggaaaatgtaagggcggaaagtaaccttatcacaaaggaatcttatcccccactacttatccttttatatttttccgtgtcatttttgcccttgagttttcctatataaggaaccaagttcggcatttgtgaaaacaagaaaaaatttggtgtaagctattttctttgaagtactgaggatacaacttcagagaaatttgtaagtttgtaatggacaagaagtactccattgggctcgatatcggcacaaacagcgtcggctgggccgtcattacggacgagtacaaggtgccgagcaaaaaattcaaagttctgggcaataccgatcgccacagcataaagaagaacctcattggcgccctcctgttcgactccggggagacggccgaagccacgcggctcaaaagaacagcacggcgcagatatacccgcagaaagaatcggatctgctacctgcaggagatctttagtaatgagatggctaaggtggatgactctttcttccataggctggaggagtcctttttggtggaggaggataaaaagcacgagcgccacccaatctttggcaatatcgtggacgaggtggcgtaccatgaaaagtacccaaccatatatcatctgaggaagaagcttgtagacagtactgataaggctgacttgcggttgatctatctcgcgctggcgcatatgatcaaatttcggggacacttcctcatcgagggggacctgaacccagacaacagcgatgtcgacaaactctttatccaactggttcagacttacaatcagcttttcgaagagaacccgatcaacgcatccggagttgacgccaaagcaatcctgagcgctaggctgtccaaatcccggcggctcgaaaacctcatcgcacagctccctggggagaagaagaacggcctgtttggtaatcttatcgccctgtcactcgggctgacccccaactttaaatctaacttcgacctggccgaagatgccaagcttcaactgagcaaagacacctacgatgatgatctcgacaatctgctggcccagatcggcgaccagtacgcagacctttttttggcggcaaagaacctgtcagacgccattctgctgagtgatattctgcgagtgaacacggagatcaccaaagctccgctgagcgctagtatgatcaagcgctatgatgagcaccaccaagacttgactttgctgaaggcccttgtcagacagcaactgcctgagaagtacaaggaaattttcttcgatcagtctaaaaatggctacgccggatacattgacggcggagcaagccaggaggaattttacaaatttattaagcccatcttggaaaaaatggacggcaccgaggagctgctggtaaagcttaacagagaagatctgttgcgcaaacagcgcactttcgacaatggaagcatcccccaccagattcacctgggcgaactgcacgctatcctcaggcggcaagaggatttctacccctttttgaaagataacagggaaaagattgagaaaatcctcacatttcggataccctactatgtaggccccctcgcccggggaaattccagattcgcgtggatgactcgcaaatcagaagagactatcactccctggaacttcgaggaagtcgtggataagggggcctctgcccagtccttcatcgaaaggatgactaactttgataaaaatctgcctaacgaaaaggtgcttcctaaacactctctgctgtacgagtacttcacagtttataacgagctcaccaaggtcaaatacgtcacagaagggatgagaaagccagcattcctgtctggagagcagaagaaagctatcgtggacctcctcttcaagacgaaccggaaagttaccgtgaaacagctcaaagaagattatttcaaaaagattgaatgtttcgactctgttgaaatcagcggagtggaggatcgcttcaacgcatccctgggaacgtatcacgatctcctgaaaatcattaaagacaaggacttcctggacaatgaggagaacgaggacattcttgaggacattgtcctcacccttacgttgtttgaagatagggagatgattgaagaacgcttgaaaacttacgctcatctcttcgacgacaaagtcatgaaacagctcaagaggcgccgatatacaggatgggggcggctgtcaagaaaactgatcaatgggatccgagacaagcagagtggaaagacaatcctggattttcttaagtccgatggatttgccaaccggaacttcatgcagttgatccatgatgactctctcacctttaaggaggacatccagaaagcacaagtttctggccagggggacagtctccacgagcacatcgctaatcttgcaggtagcccagctatcaaaaagggaatactgcagaccgttaaggtcgtggatgaactcgtcaaagtaatgggaaggcataagcccgagaatatcgttatcgagatggcccgagagaaccaaactacccagaagggacagaagaacagtagggaaaggatgaagaggattgaagagggtataaaagaactggggtcccaaatccttaaggaacacccagttgaaaacacccagcttcagaatgagaagctctacctgtactacctgcagaacggcagggacatgtacgtggatcaggaactggacatcaatcggctctccgactacgacgtggatcatatcgtgccccagtcttttctcaaagatgattctattgataataaagtgttgacaagatccgataaaaatagagggaagagtgataacgtcccctcagaagaagttgtcaagaaaatgaaaaattattggcggcagctgctgaacgccaaactgatcacacaacggaagttcgataatctgactaaggctgaacgaggtggcctgtctgagttggataaagccggcttcatcaaaaggcagcttgttgagacacgccagatcaccaagcacgtggcccaaattctcgattcacgcatgaacaccaagtacgatgaaaatgacaaactgattcgagaggtgaaagttattactctgaagtctaagctggtttcagatttcagaaaggactttcagttttataaggtgagagagatcaacaattaccaccatgcgcatgatgcctacctgaatgcagtggtaggcactgcacttatcaaaaaatatcccaagcttgaatctgaatttgtttacggagactataaagtgtacgatgttaggaaaatgatcgcaaagtctgagcaggaaataggcaaggccaccgctaagtacttcttttacagcaatattatgaattttttcaagaccgagattacactggccaatggagagattcggaagcgaccacttatcgaaacaaacggagaaacaggagaaatcgtgtgggacaagggtagggatttcgcgacagtccggaaggtcctgtccatgccgcaggtgaacatcgttaaaaagaccgaagtacagaccggaggcttctccaaggaaagtatcctcccgaaaaggaacagcgacaagctgatcgcacgcaaaaaagattgggaccccaagaaatacggcggattcgattctcctacagtcgcttacagtgtactggttgtggccaaagtggagaaagggaagtctaaaaaactcaaaagcgtcaaggaactgctgggcatcacaatcatggagcgatcaagcttcgaaaaaaaccccatcgactttctcgaggcgaaaggatataaagaggtcaaaaaagacctcatcattaagcttcccaagtactctctctttgagcttgaaaacggccggaaacgaatgctcgctagtgcgggcgagctgcagaaaggtaacgagctggcactgccctctaaatacgttaatttcttgtatctggccagccactatgaaaagctcaaaggatctcccgaagataatgagcagaagcagctgttcgtggaacaacacaaacactaccttgatgagatcatcgagcaaataagcgaattctccaaaagagtgatcctcgccgacgctaacctcgataaggtgctttctgcttacaataagcacagggataagcccatcagggagcaggcagaaaacattatccacttgtttactctgaccaacttgggcgcgcctgcagccttcaagtacttcgacaccaccatagacagaaagcggtacacctctacaaaggaggtcctggacgccacactgattcatcagtcaattacggggctctatgaaacaagaatcgacctctctcagctcggtggagacagcagggctgaccccaagaagaagaggaaggtgtgagcttctctagctagagtcgatcgacaagctcgagtttctccataataatgtgtgagtagttcccagataagggaattagggttcctatagggtttcgctcatgtgttgagcatataagaaacccttagtatgtatttgtatttgtaaaatacttctatcaataaaatttctaattcctaaaaccaaaatccagtactaaaatccagat

**S1 guide cassette:**

*At* U626 promoter

Variable protospacer sequence

Invariable sgRNA sequence

Terminator

catcttcattcttaagatatgaagataatcttcaaaaggcccctgggaatctgaaagaagagaagcaggcccatttatatgggaaagaacaatagtatttcttatataggcccatttaagttgaaaacaatcttcaaaagtcccacatcgcttagataagaaaacgaagctgagtttatatacagctagagtcgaagtagtgattgNNNNNNNNNNNNNNNNNNNNgttttagagctagaaatagcaagttaaaataaggctagtccgttatcaacttgaaaaagtggcaccgagtcggtgctttttttctagacccagctttcttgtacaaagttggcattacgct

**S2 *At*CYS4-Cas9 cassette:**

*At*Ubi10 promoter

Cys4 CDS

P2A self cleaving peptide CDS

*At*Cas9 CDS

*Pisum sativum* E9 terminator

GTCGAGCTGCAGGTCAACGGATCAGGATATTCTTGTTTAAGATGTTGAACTCTATGGAGGTTTGTATGAACTGATGATCTAGGACCGGATAAGTTCCCTTCTTCATAGCGAACTTATTCAAAGAATGTTTTGTGTATCATTCTTGTTACATTGTTATTAATGAAAAAATATTATTGGTCATTGGACTGAACACGAGTGTTAAATATGGACCAGGCCCCAAATAAGATCCATTGATATATGAATTAAATAACAAGAATAAATCGAGTCACCAAACCACTTGCCTTTTTTAACGAGACTTGTTCACCAACTTGATACAAAAGTCATTATCCTATGCAAATCAATAATCATACAAAAATATCCAATAACACTAAAAAATTAAAAGAAATGGATAATTTCACAATATGTTATACGATAAAGAAGTTACTTTTCCAAGAAATTCACTGATTTTATAAGCCCACTTGCATTAGATAAATGGCAAAAAAAAACAAAAAGGAAAAGAAATAAAGCACGAAGAATTCTAGAAAATACGAAATACGCTTCAATGCAGTGGGACCCACGGTTCAATTATTGCCAATTTTCAGCTCCACCGTATATTTAAAAAATAAAACGATAATGCTAAAAAAATATAAATCGTAACGATCGTTAAATCTCAACGGCTGGATCTTATGACGACCGTTAGAAATTGTGGTTGTCGACGAGTCAGTAATAAACGGCGTCAAAGTGGTTGCAGCCGGCACACACGAGTCGTGTTTATCAACTCAAAGCACAAATACTTTTCCTCAACCTAAAAATAAGGCAATTAGCCAAAAACAACTTTGCGTGTAAACAACGCTCAATACACGTGTCATTTTATTATTAGCTATTGCTTCACCGCCTTAGCTTTCTCGTGACCTAGTCGTCCTCGTCTTTTCTTCTTCTTCTTCTATAAAACAATACCCAAAGAGCTCTTCTTCTTCACAATTCAGATTTCAATTTCTCAAAATCTTAAAAACTTTCTCTCAATTCTCTCTACCGTGATCAAGGTAAATTTCTGTGTTCCTTATTCTCTCAAAATCTTCGATTTTGTTTTCGTTCGATCCCAATTTCGTATATGTTCTTTGGTTTAGATTCTGTTAATCTTAGATCGAACACGATTTTCTGGGTTTGATCGTTAGATATCATCTTAATTCTCGATTAGGGTTTCATAGATATCATCCGATTTGTTCAAATAATTTGAGTTTTGTCGAATAATTACTCTTCGATTTGTGATTTCTATCTAGATCTGGTGTTAGTTTCTAGTTTGTGCGATCGAATTTGTCGATTAATCTGAGTTTTTCTGATTAACAGGAATGgatcattatcttgatattagacttagacctgatccagaatttccaccagctcaacttatgtctgttctttttggaaaacttcatcaagctcttgttgctcaaggaggagatagaattggagtttcttttcctgatcttgatgaatcaagatcaagacttggagaaagacttagaattcatgcttctgctgatgatcttagagctttgcttgctagaccttggcttgaaggacttagagatcatcttcaatttggagaaccagctgttgttccacatccaactccttatagacaagtttcaagagttcaagctaaatctaatccagaaagacttagaaggagacttatgaggagacatgatctttctgaagaagaagctagaaaaagaattcctgatactgttgctagagctttggatttgccttttgttacacttagatcacaatctactggacaacattttagactttttattagacatggaccacttcaagttactgctgaagaaggaggatttacttgttatggactttctaagggaggttttgttccttggtttggatctggagctactaatttttctcttcttaagcaagctggagatgttgaagaaaatcctggacccatggataagaagtactctatcggactcgatatcggaactaactctgtgggatgggctgtgatcaccgatgagtacaaggtgccatctaagaagttcaaggttctcggaaacaccgataggcactctatcaagaaaaaccttatcggtgctctcctcttcgattctggtgaaactgctgaggctaccagactcaagagaaccgctagaagaaggtacaccagaagaaagaacaggatctgctacctccaagagatcttctctaacgagatggctaaagtggatgattcattcttccacaggctcgaagagtcattcctcgtggaagaagataagaagcacgagaggcaccctatcttcggaaacatcgttgatgaggtggcataccacgagaagtaccctactatctaccacctcagaaagaagctcgttgattctactgataaggctgatctcaggctcatctacctcgctctcgctcacatgatcaagttcagaggacacttcctcatcgagggtgatctcaaccctgataactctgatgtggataagttgttcatccagctcgtgcagacctacaaccagcttttcgaagagaaccctatcaacgcttcaggtgtggatgctaaggctatcctctctgctaggctctctaagtcaagaaggcttgagaacctcattgctcagctccctggtgagaagaagaacggacttttcggaaacttgatcgctctctctctcggactcacccctaacttcaagtctaacttcgatctcgctgaggatgcaaagctccagctctcaaaggatacctacgatgatgatctcgataacctcctcgctcagatcggagatcagtacgctgatttgttcctcgctgctaagaacctctctgatgctatcctcctcagtgatatcctcagagtgaacaccgagatcaccaaggctccactctcagcttctatgatcaagagatacgatgagcaccaccaggatctcacacttctcaaggctcttgttagacagcagctcccagagaagtacaaagagattttcttcgatcagtctaagaacggatacgctggttacatcgatggtggtgcatctcaagaagagttctacaagttcatcaagcctatcctcgagaagatggatggaaccgaggaactcctcgtgaagctcaatagagaggatcttctcagaaagcagaggaccttcgataacggatctatccctcatcagatccacctcggagagttgcacgctatccttagaaggcaagaggatttctacccattcctcaaggataacagggaaaagattgagaagattctcaccttcagaatcccttactacgtgggacctctcgctagaggaaactcaagattcgcttggatgaccagaaagtctgaggaaaccatcaccccttggaacttcgaagaggtggtggataagggtgctagtgctcagtctttcatcgagaggatgaccaacttcgataagaaccttccaaacgagaaggtgctccctaagcactctttgctctacgagtacttcaccgtgtacaacgagttgaccaaggttaagtacgtgaccgagggaatgaggaagcctgcttttttgtcaggtgagcaaaagaaggctatcgttgatctcttgttcaagaccaacagaaaggtgaccgtgaagcagctcaaagaggattacttcaagaaaatcgagtgcttcgattcagttgagatttctggtgttgaggataggttcaacgcatctctcggaacctaccacgatctcctcaagatcattaaggataaggatttcttggataacgaggaaaacgaggatatcttggaggatatcgttcttaccctcaccctctttgaagatagagagatgattgaagaaaggctcaagacctacgctcatctcttcgatgataaggtgatgaagcagttgaagagaagaagatacactggttggggaaggctctcaagaaagctcattaacggaatcagggataagcagtctggaaagacaatccttgatttcctcaagtctgatggattcgctaacagaaacttcatgcagctcatccacgatgattctctcacctttaaagaggatatccagaaggctcaggtttcaggacagggtgatagtctccatgagcatatcgctaacctcgctggatctcctgcaatcaagaagggaatcctccagactgtgaaggttgtggatgagttggtgaaggtgatgggaaggcataagcctgagaacatcgtgatcgaaatggctagagagaaccagaccactcagaagggacagaagaactctagggaaaggatgaagaggatcgaggaaggtatcaaagagcttggatctcagatcctcaaagagcaccctgttgagaacactcagctccagaatgagaagctctacctctactacctccagaacggaagggatatgtatgtggatcaagagttggatatcaacaggctctctgattacgatgttgatcatatcgtgccacagtcattcttgaaggatgattctatcgataacaaggtgctcaccaggtctgataagaacaggggtaagagtgataacgtgccaagtgaagaggttgtgaagaaaatgaagaactattggaggcagctcctcaacgctaagctcatcactcagagaaagttcgataacttgactaaggctgagaggggaggactctctgaattggataaggcaggattcatcaagaggcagcttgtggaaaccaggcagatcactaagcacgttgcacagatcctcgattctaggatgaacaccaagtacgatgagaacgataagttgatcagggaagtgaaggttatcaccctcaagtcaaagctcgtgtctgatttcagaaaggatttccaattctacaaggtgagggaaatcaacaactaccaccacgctcacgatgcttaccttaacgctgttgttggaaccgctctcatcaagaagtatcctaagctcgagtcagagttcgtgtacggtgattacaaggtgtacgatgtgaggaagatgatcgctaagtctgagcaagagatcggaaaggctaccgctaagtatttcttctactctaacatcatgaatttcttcaagaccgagattaccctcgctaacggtgagatcagaaagaggccactcatcgagacaaacggtgaaacaggtgagatcgtgtgggataagggaagggatttcgctaccgttagaaaggtgctctctatgccacaggtgaacatcgttaagaaaaccgaggtgcagaccggtggattctctaaagagtctatcctccctaagaggaactctgataagctcattgctaggaagaaggattgggaccctaagaaatacggtggtttcgattctcctaccgtggcttactctgttctcgttgtggctaaggttgagaagggaaagagtaagaagctcaagtctgttaaggaacttctcggaatcactatcatggaaaggtcatctttcgagaagaacccaatcgatttcctcgaggctaagggatacaaagaggttaagaaggatctcatcatcaagctcccaaagtactcactcttcgaactcgagaacggtagaaagaggatgctcgcttctgctggtgagcttcaaaagggaaacgagcttgctctcccatctaagtacgttaactttctttacctcgcttctcactacgagaagttgaagggatctccagaagataacgagcagaagcaacttttcgttgagcagcacaagcactacttggatgagatcatcgagcagatctctgagttctctaaaagggtgatcctcgctgatgcaaacctcgataaggtgttgtctgcttacaacaagcacagagataagcctatcagggaacaggcagagaacatcatccatctcttcacccttaccaacctcggtgctcctgctgctttcaagtacttcgatacaaccatcgataggaagagatacacctctaccaaagaagtgctcgatgctaccctcatccatcagtctatcactggactctacgagactaggatcgatctctcacagctcggtggtgattcaagggctgatcctaagaagaagaggaaggtttgagcttgctttcgttcgtatcatcggtttcgacaacgttcgtcaagttcaatgcatcagtttcattgcgcacacaccagaatcctactgagtttgagtattatggcattgggaaaactgtttttcttgtaccatttgttgtgcttgtaatttactgtgttttttattcggttttcgctatcgaactgtgaaatggaaatggatggagaagagttaatgaatgatatggtccttttgttcattctcaaattaatattatttgttttttctcttatttgttgtgtgttgaatttgaaattataagagatatgcaaacattttgttttgagtaaaaatgtgtcaaatcgtggcctctaatgaccgaagttaatatgaggagtaaaacacttgtagttgtaccattatgcttattcactaggcaacaaatatattttcagacctagaaaagctgcaaatgttactgaatacaagtatgtcctcttgtgttttagacatttatgaactttcctttatgtaattttccagaatccttgtcagattctaatcattgctttataattatagttatactcatggatttgtagttgagtatgaaaatattttttaatgcattttatgacttgccaattgattgacaac

**S2 guide cassette:**

*At*Ubi5 promoter

Cys4 cleavage site

Variable spacer sequence

Invariable sgRNA sequence

35s terminator

cgggttgatcaatatctttcccttgttctgatcctatgtaactacatcttttgttccgagaaaattgattttatagtttttttgttttcttttcttttttcttgattttatagagttattacatatatttaaaaataattcttgtttaattatttaataattttcattttatatgtgaaatatttttttataattttaaaccaatgaatttttagatatttttactttttaatgacattatgatatttatgtttttaaaaaaaaaattaacgatgaaaagttaatcctataatttatattgatgttatacttaatttatacaaaaattgaaaggtttaaataaaatcttgtttttttttaaaacgtcgtcgatctgttttgaaaacgttgcaatttttgttaatccaatttatggaaacatgttaaagcgttttaaatcaacgtttaaattttctttaatgttagctaaatgagattttttttttcaaacttacgaattattacttttaaacctgtgatacggttcaaagtaagtggatatacacaagaagtcaaatagctcccgtattttcatacaacaacaccatctttcttgaaaaaatagaccatatttttcgaggtgatctctcgttagtatccgtcaagctttcatctttatgttagcactttttttcttggttcaacttcaattttcctctagattgttctagttgttttgatttgtctcaagtcgatgttaaagtttacatgttttcctatccggagttgatatggcggtgaatgatagttggttcgatttgatggtgttcttgttcgtgagacagatttttttcgagtagttaagcttcgctccttgtctttgatttgcatatttggtagtgatttgatagcgttttgtagaattttatttagacgtaaattggtggcatatttattgtaattttacatatcagattatgagtttgaatggtaacacaatacagctttatataattttgtacaactttcacagttaccaataacaaaaactttattaaaagtgttataaaaagtttactctcagcttttttgtacaactctttctaacagctttgcactattcaaataaagctaacagctaaaaaacttgtacagcttttggttaccaatcagacactatgtaaatttttggtttttgcttttgttacttttgagaggtcggctcttttcttgataggataaaattgttgttatttttgagtttgctagcatcaataatcacaattaagatgtatcttattttgagaggtcggctctttctttgataggataaaattgttgttatttttaagtttgctgagagaattcaagatgataaaaaaattaaaaaaatcacaatcacaaggttttaataataaggtttcaaatataaacttttttttttttgcacaaactttcaaatataaacaaaaaaataattttaatacaaagtctatatatggagaaaataggaagccaaaattgataattcacaaaattaaagtaaaatctattagccgacaaaaaaaaaggtaaaatctattaaaatatagataaggttctagaaattaaataaatctattaccaattctcaaaccgacataagtacgaccaaaaaaaagtttataaataaaagtcacaacgagccttaacgcgtagaatcttcccgtactttacttttccggaggaatagaaaattgggggctagggttcgcaattgtagttttcgagcgaagaagttcactgccgtataggcagNNNNNNNNNNNNNNNNNNNNgttttagagctagaaatagcaagttaaaataaggctagtccgttatcaacttgaaaaagtggcaccgagtcggtgcgttcactgccgtataggcagNNNNNNNNNNNNNNNNNNNNgttttagagctagaaatagcaagttaaaataaggctagtccgttatcaacttgaaaaagtggcaccgagtcggtgcgttcactgccgtataggcagNNNNNNNNNNNNNNNNNNNNgttttagagctagaaatagcaagttaaaataaggctagtccgttatcaacttgaaaaagtggcaccgagtcggtgcgttcactgccgtataggcagNNNNNNNNNNNNNNNNNNNNgttcactgccgtataggcagctcgagtttctccataataatgtgtgagtagttcccagataagggaattagggttcctatagggtttcgctcatgtgttgagcatataagaaacccttagtatgtatttgtatttgtaaaatacttctatcaataaaatttctaattcctaaaaccaaaatccagtactaaaatccagatcccccgaatta

**S3 *At*Cas9 with 1 intron cassette**

*At*Ubi10 promoter

PCas9 coding sequence with one intron

Intron

*Pisum sativum* E9 terminator

GTCGAGCTGCAGGTCAACGGATCAGGATATTCTTGTTTAAGATGTTGAACTCTATGGAGGTTTGTATGAACTGATGATCTAGGACCGGATAAGTTCCCTTCTTCATAGCGAACTTATTCAAAGAATGTTTTGTGTATCATTCTTGTTACATTGTTATTAATGAAAAAATATTATTGGTCATTGGACTGAACACGAGTGTTAAATATGGACCAGGCCCCAAATAAGATCCATTGATATATGAATTAAATAACAAGAATAAATCGAGTCACCAAACCACTTGCCTTTTTTAACGAGACTTGTTCACCAACTTGATACAAAAGTCATTATCCTATGCAAATCAATAATCATACAAAAATATCCAATAACACTAAAAAATTAAAAGAAATGGATAATTTCACAATATGTTATACGATAAAGAAGTTACTTTTCCAAGAAATTCACTGATTTTATAAGCCCACTTGCATTAGATAAATGGCAAAAAAAAACAAAAAGGAAAAGAAATAAAGCACGAAGAATTCTAGAAAATACGAAATACGCTTCAATGCAGTGGGACCCACGGTTCAATTATTGCCAATTTTCAGCTCCACCGTATATTTAAAAAATAAAACGATAATGCTAAAAAAATATAAATCGTAACGATCGTTAAATCTCAACGGCTGGATCTTATGACGACCGTTAGAAATTGTGGTTGTCGACGAGTCAGTAATAAACGGCGTCAAAGTGGTTGCAGCCGGCACACACGAGTCGTGTTTATCAACTCAAAGCACAAATACTTTTCCTCAACCTAAAAATAAGGCAATTAGCCAAAAACAACTTTGCGTGTAAACAACGCTCAATACACGTGTCATTTTATTATTAGCTATTGCTTCACCGCCTTAGCTTTCTCGTGACCTAGTCGTCCTCGTCTTTTCTTCTTCTTCTTCTATAAAACAATACCCAAAGAGCTCTTCTTCTTCACAATTCAGATTTCAATTTCTCAAAATCTTAAAAACTTTCTCTCAATTCTCTCTACCGTGATCAAGGTAAATTTCTGTGTTCCTTATTCTCTCAAAATCTTCGATTTTGTTTTCGTTCGATCCCAATTTCGTATATGTTCTTTGGTTTAGATTCTGTTAATCTTAGATCGAACACGATTTTCTGGGTTTGATCGTTAGATATCATCTTAATTCTCGATTAGGGTTTCATAGATATCATCCGATTTGTTCAAATAATTTGAGTTTTGTCGAATAATTACTCTTCGATTTGTGATTTCTATCTAGATCTGGTGTTAGTTTCTAGTTTGTGCGATCGAATTTGTCGATTAATCTGAGTTTTTCTGATTAACAGGAATGGATTACAAGGATGATGATGATAAGGATTACAAGGATGATGATGATAAGATGGCTCCAAAGAAGAAGAGAAAGGTTGGAATCCACGGAGTTCCAGCTGCTGGAGGTATGGATAAGAAGTACTCTATCGGACTTGACATCGGAACCAACTCTGTTGGATGGGCTGTTATCACCGATGAGTACAAGGTTCCATCTAAGAAGTTCAAGGTTCTTGGAAACACCGATAGACACTCTATCAAGAAGAACCTTATCGGTGCTCTTCTTTTCGATTCTGGAGAAACCGCTGAGGCTACCAGATTGAAGAGAACCGCTAGAAGAAGATACACCAGAAGAAAGAACAGAATCTGCTACCTTCAGGAAATCTTCTCTAACGAGATGGCTAAGGTTGATGATTCTTTCTTCCACAGACTTGAGGAGTCTTTCCTTGTTGAGGAGGATAAGAAGCACGAGAGACACCCAATCTTCGGAAACATCGTTGATGAGGTTGCTTACCACGAGAAGTACCCAACCATCTACCACCTTAGAAAGAAGTTGGTTGATTCTACCGATAAGGCTGATCTTAGACTTATCTACCTTGCTCTTGCTCACATGATCAAGTTCAGAGGACACTTCCTTATCGAGGGAGATCTTAACCCAGATAACTCTGATGTTGATAAGTTGTTCATCCAGCTTGTTCAGACCTACAACCAGCTTTTCGAGGAGAACCCAATCAACGCTTCTGGAGTTGATGCTAAGGCTATCCTTTCTGCTAGACTTTCTAAGTCTCGTAGACTTGAGAACCTTATCGCTCAGCTTCCAGGAGAGAAGAAGAACGGACTTTTCGGAAACCTTATCGCTCTTTCTCTTGGACTTACCCCAAACTTCAAGTCTAACTTCGATCTTGCTGAGGATGCTAAGTTGCAGCTTTCTAAGGATACCTACGATGATGATCTTGATAACCTTCTTGCTCAGATCGGAGATCAGTACGCTGATCTTTTCCTTGCTGCTAAGAACCTTTCTGATGCTATCCTTCTTTCTGACATCCTTAGAGTTAACACCGAGATCACCAAGGCTCCACTTTCTGCTTCTATGATCAAGAGATACGATGAGCACCACCAGGATCTTACCCTTTTGAAGGCTCTTGTTAGACAGCAGCTTCCAGAGAAGTACAAGGAAATCTTCTTCGATCAGTCTAAGAACGGATACGCTGGATACATCGATGGAGGAGCTTCTCAGGAGGAGTTCTACAAGTTCATCAAGCCAATCCTTGAGAAGATGGATGGAACCGAGGAGCTTCTTGTTAAGTTGAACAGAGAGGATCTTCTTAGAAAGCAGAGAACCTTCGATAACGGATCTATCCCACACCAGATCCACCTTGGAGAGCTTCACGCTATCCTTCGTAGACAGGAGGATTTCTACCCATTCTTGAAGGATAACAGAGAGAAGATCGAGAAGATCCTTACCTTCAGAATCCCATACTACGTTGGACCACTTGCTAGAGGAAACTCTCGTTTCGCTTGGATGACCAGAAAGTCTGAGGAGACAATCACCCCTTGGAACTTCGAGGAGGTAAGTTTCTGCTTCTACCTTTGATATATATATAATAATTATCATTAATTAGTAGTAATATAATATTTCAAATATTTTTTTCAAAATAAAAGAATGTAGTATATAGCAATTGCTTTTCTGTAGTTTATAAGTGTGTATATTTTAATTTATAACTTTTCTAATATATGACCAAAATTTGTTGATGTGCAGGTTGTTGATAAGGGAGCTTCTGCTCAGTCTTTCATCGAGAGAATGACCAACTTCGATAAGAACCTTCCAAACGAGAAGGTTCTTCCAAAGCACTCTCTTCTTTACGAGTACTTCACCGTTTACAACGAGCTTACCAAGGTTAAGTACGTTACCGAGGGAATGAGAAAGCCAGCTTTCCTTTCTGGAGAGCAGAAGAAGGCTATCGTTGATCTTCTTTTCAAGACCAACAGAAAGGTTACCGTTAAGCAGTTGAAGGAGGATTACTTCAAGAAGATCGAGTGCTTCGATTCTGTTGAAATCTCTGGAGTTGAGGATAGATTCAACGCTTCTCTTGGAACCTACCACGATCTTTTGAAGATCATCAAGGATAAGGATTTCCTTGATAACGAGGAGAACGAGGACATCCTTGAGGACATCGTTCTTACCCTTACCCTTTTCGAGGATAGAGAGATGATCGAGGAGAGACTCAAGACCTACGCTCACCTTTTCGATGATAAGGTTATGAAGCAGTTGAAGAGAAGAAGATACACCGGATGGGGTAGACTTTCTCGTAAGTTGATCAACGGAATCAGAGATAAGCAGTCTGGAAAGACCATCCTTGATTTCTTGAAGTCTGATGGATTCGCTAACAGAAACTTCATGCAGCTTATCCACGATGATTCTCTTACCTTCAAGGAGGACATCCAGAAGGCTCAGGTTTCTGGACAGGGAGATTCTCTTCACGAGCACATCGCTAACCTTGCTGGATCTCCAGCTATCAAGAAGGGAATCCTTCAGACCGTTAAGGTTGTTGATGAGCTTGTTAAGGTTATGGGTAGACACAAGCCAGAGAACATCGTTATCGAGATGGCTAGAGAGAACCAGACCACCCAGAAGGGACAGAAGAACTCTCGTGAGAGAATGAAGAGAATCGAGGAGGGAATCAAGGAGCTTGGATCTCAAATCTTGAAGGAGCACCCAGTTGAGAACACCCAGCTTCAGAACGAGAAGTTGTACCTTTACTACCTTCAGAACGGAAGAGATATGTACGTTGATCAGGAGCTTGACATCAACAGACTTTCTGATTACGATGTTGATCACATCGTTCCACAGTCTTTCTTGAAGGATGATTCTATCGATAACAAGGTTCTTACCCGTTCTGATAAGAACAGAGGAAAGTCTGATAACGTTCCATCTGAGGAGGTTGTTAAGAAGATGAAGAACTACTGGAGACAGCTTCTTAACGCTAAGTTGATCACCCAGAGAAAGTTCGATAACCTTACCAAGGCTGAGAGAGGAGGACTTTCTGAGCTTGATAAGGCTGGATTCATCAAGAGACAGCTTGTTGAAACCAGACAGATCACCAAGCACGTTGCTCAGATCCTTGATTCTCGTATGAACACCAAGTACGATGAGAACGATAAGTTGATCAGAGAGGTTAAGGTTATCACCTTGAAGTCTAAGTTGGTTTCTGATTTCAGAAAGGATTTCCAGTTCTACAAGGTTAGAGAGATCAACAACTACCACCACGCTCACGATGCTTACCTTAACGCTGTTGTTGGAACCGCTCTTATCAAGAAGTACCCAAAGTTGGAGTCTGAGTTCGTTTACGGAGATTACAAGGTTTACGATGTTAGAAAGATGATCGCTAAGTCTGAGCAGGAGATCGGAAAGGCTACCGCTAAGTACTTCTTCTACTCTAACATCATGAACTTCTTCAAGACCGAGATCACCCTTGCTAACGGAGAGATCAGAAAAAGACCACTTATCGAGACAAACGGAGAGACAGGAGAGATCGTTTGGGATAAGGGAAGAGATTTCGCTACCGTTAGAAAGGTTCTTTCTATGCCACAGGTTAACATCGTTAAGAAAACCGAGGTTCAGACCGGAGGATTCTCTAAGGAGTCTATCCTTCCAAAGAGAAACTCTGATAAGTTGATCGCTAGAAAGAAGGATTGGGACCCAAAGAAGTACGGAGGATTCGATTCTCCAACCGTTGCTTACTCTGTTCTTGTTGTTGCTAAGGTTGAGAAGGGAAAGTCTAAGAAGTTGAAGTCTGTTAAGGAGCTTCTTGGAATCACCATCATGGAGCGTTCTTCTTTCGAGAAGAACCCAATCGATTTCCTTGAGGCTAAGGGATACAAGGAGGTTAAGAAGGATCTTATCATCAAGTTGCCAAAGTACTCTCTTTTCGAGCTTGAGAACGGAAGAAAGAGAATGCTTGCTTCTGCTGGAGAGCTTCAGAAGGGAAACGAGCTTGCTCTTCCATCTAAGTACGTTAACTTCCTTTACCTTGCTTCTCACTACGAGAAGTTGAAGGGATCTCCAGAGGATAACGAGCAGAAGCAGCTTTTCGTTGAGCAGCACAAGCACTACCTTGATGAGATCATCGAGCAAATCTCTGAGTTCTCTAAGAGAGTTATCCTTGCTGATGCTAACCTTGATAAGGTTCTTTCTGCTTACAACAAGCACAGAGATAAGCCAATCAGAGAGCAGGCTGAGAACATCATCCACCTTTTCACCCTTACCAACCTTGGTGCTCCAGCTGCTTTCAAGTACTTCGATACCACCATCGATAGAAAAAGATACACCTCTACCAAGGAGGTTCTTGATGCTACCCTTATCCACCAGTCTATCACCGGACTTTACGAGACAAGAATCGATCTTTCTCAGCTTGGAGGAGATGGTGGCAAGAGGCCAGCTGCTACCAAGAAGGCTGGACAGGCTAAGAAGAAGAAGTGAGCTTGCTTTCGTTCGTATCATCGGTTTCGACAACGTTCGTCAAGTTCAATGCATCAGTTTCATTGCGCACACACCAGAATCCTACTGAGTTTGAGTATTATGGCATTGGGAAAACTGTTTTTCTTGTACCATTTGTTGTGCTTGTAATTTACTGTGTTTTTTATTCGGTTTTCGCTATCGAACTGTGAAATGGAAATGGATGGAGAAGAGTTAATGAATGATATGGTCCTTTTGTTCATTCTCAAATTAATATTATTTGTTTTTTCTCTTATTTGTTGTGTGTTGAATTTGAAATTATAAGAGATATGCAAACATTTTGTTTTGAGTAAAAATGTGTCAAATCGTGGCCTCTAATGACCGAAGTTAATATGAGGAGTAAAACACTTGTAGTTGTACCATTATGCTTATTCACTAGGCAACAAATATATTTTCAGACCTAGAAAAGCTGCAAATGTTACTGAATACAAGTATGTCCTCTTGTGTTTTAGACATTTATGAACTTTCCTTTATGTAATTTTCCAGAATCCTTGTCAGATTCTAATCATTGCTTTATAATTATAGTTATACTCATGGATTTGTAGTTGAGTATGAAAATATTTTTTAATGCATTTTATGACTTGCCAATTGATTGACAA

**S3 guide cassette**

*At*U626 promoter

tRNA sequence

Variable spacer sequence

Invariable sgRNA sequence

Terminator

CatcttcattcttaagatatgaagataatcttcaaaaggcccctgggaatctgaaagaagagaagcaggcccatttatatgggaaagaacaatagtatttcttatataggcccatttaagttgaaaacaatcttcaaaagtcccacatcgcttagataagaaaacgaagctgagtttatatacagctagagtcgaagtagtgattgaacaaagcaccagtggtctagtggtagaatagtaccctgccacggtacagacccgggttcgattcccggctggtgcaNNNNNNNNNNNNNNNNNNNNgttttagagctagaaatagcaagttaaaataaggctagtccgttatcaacttgaaaaagtggcaccgagtcggtgcaacaaagcaccagtggtctagtggtagaatagtaccctgccacggtacagacccgggttcgattcccggctggtgcaNNNNNNNNNNNNNNNNNNNNgttttagagctagaaatagcaagttaaaataaggctagtccgttatcaacttgaaaaagtggcaccgagtcggtgcaacaaagcaccagtggtctagtggtagaatagtaccctgccacggtacagacccgggttcgattcccggctggtgcaNNNNNNNNNNNNNNNNNNNNgttttagagctagaaatagcaagttaaaataaggctagtccgttatcaacttgaaaaagtggcaccgagtcggtgcaacaaagcaccagtggtctagtggtagaatagtaccctgccacggtacagacccgggttcgattcccggctggtgcaNNNNNNNNNNNNNNNNNNNNgttttagagctagaaatagcaagttaaaataaggctagtccgttatcaacttgaaaaagtggcaccgagtcggtgcttttttt

**S4 *At*Cas9 cassette with 13 introns cassette**

*At*Ubi10 promoter

ZCas9 CDS with 13 introns

Intron

*Pisum sativum* E9 terminator

GGAGGTCGAGCTGCAGGTCAACGGATCAGGATATTCTTGTTTAAGATGTTGAACTCTATGGAGGTTTGTATGAACTGATGATCTAGGACCGGATAAGTTCCCTTCTTCATAGCGAACTTATTCAAAGAATGTTTTGTGTATCATTCTTGTTACATTGTTATTAATGAAAAAATATTATTGGTCATTGGACTGAACACGAGTGTTAAATATGGACCAGGCCCCAAATAAGATCCATTGATATATGAATTAAATAACAAGAATAAATCGAGTCACCAAACCACTTGCCTTTTTTAACGAGACTTGTTCACCAACTTGATACAAAAGTCATTATCCTATGCAAATCAATAATCATACAAAAATATCCAATAACACTAAAAAATTAAAAGAAATGGATAATTTCACAATATGTTATACGATAAAGAAGTTACTTTTCCAAGAAATTCACTGATTTTATAAGCCCACTTGCATTAGATAAATGGCAAAAAAAAACAAAAAGGAAAAGAAATAAAGCACGAAGAATTCTAGAAAATACGAAATACGCTTCAATGCAGTGGGACCCACGGTTCAATTATTGCCAATTTTCAGCTCCACCGTATATTTAAAAAATAAAACGATAATGCTAAAAAAATATAAATCGTAACGATCGTTAAATCTCAACGGCTGGATCTTATGACGACCGTTAGAAATTGTGGTTGTCGACGAGTCAGTAATAAACGGCGTCAAAGTGGTTGCAGCCGGCACACACGAGTCGTGTTTATCAACTCAAAGCACAAATACTTTTCCTCAACCTAAAAATAAGGCAATTAGCCAAAAACAACTTTGCGTGTAAACAACGCTCAATACACGTGTCATTTTATTATTAGCTATTGCTTCACCGCCTTAGCTTTCTCGTGACCTAGTCGTCCTCGTCTTTTCTTCTTCTTCTTCTATAAAACAATACCCAAAGAGCTCTTCTTCTTCACAATTCAGATTTCAATTTCTCAAAATCTTAAAAACTTTCTCTCAATTCTCTCTACCGTGATCAAGGTAAATTTCTGTGTTCCTTATTCTCTCAAAATCTTCGATTTTGTTTTCGTTCGATCCCAATTTCGTATATGTTCTTTGGTTTAGATTCTGTTAATCTTAGATCGAACACGATTTTCTGGGTTTGATCGTTAGATATCATCTTAATTCTCGATTAGGGTTTCATAGATATCATCCGATTTGTTCAAATAATTTGAGTTTTGTCGAATAATTACTCTTCGATTTGTGATTTCTATCTAGATCTGGTGTTAGTTTCTAGTTTGTGCGATCGAATTTGTCGATTAATCTGAGTTTTTCTGATTAACAGGaatggcttctagcccaccgaagaagaagcggaaggtcagctggaaaatggacaagaagtacagcattggacttgatattggtacgaactcagttgggtgggccgttatcaccgatgaatacaaggtaccttcgaagaaatttaaagtgctgggcaacacagataggcacagcattaagaagaacttgatcggagctctgctctttgactctggagaaaccgcggaggcgacaaggcttaaacgtactgcgaggagaaggtacactcgcaggaagaacagaatctgttatctccaagagatctttagcaacgagatggcgaaggtaaggatttttatgatatagtatgcttatgtattttgtactgaaagcatatcctgcttcattgggatattactgaaagcatttaactacatgtaaactcacttgatgatcaataaacttgattttgcaggttgacgactcgttcttccatcgcctcgaggaatccttcctggtagaggaagataagaaacacgagcgtcaccccatctttgggaatattgttgacgaagtagcctatcatgaaaagtatccgactatataccaccttcgcaagaagctggtggactcaaccgataaggcagaccttcggctcatatacctggctctcgcgcacatgataaagtttcgtggccatttcttgatcgaaggggacctcaacccggataactccgatgtggataaactgttcattcagctcgtccaaacctacaatcagctgttcgaggagaaccccatcaatgcatcaggtaacattccttagttacctttcttttctttttccatcataagtttatagattgtacatgctttgagatttttctttgcaaacaatctcaggtgtcgacgccaaggcaatactgtctgccagactttcgaagtccagacggcttgagaatctgatcgctcaattgccaggcgagaagaagaacggcttgttcgggaatctgattgcactgtctctgggcctcacccctaacttcaaaagcaactttgacctcgccgaggacgcgaagctgcagctgtcaaaggatacatacgatgatgatctggacaatctgctcgcccaaataggtgctcttgaaattggaactcttcttttgttgtctaaacctatcaatttctttgcggaaatttatttgaagctgtagagttaaaattgagtcttttaaacttttgtaggtgatcagtatgccgacctgttcttggctgccaagaatctgtcagacgctatcttgctcagtgacattctgcgggtcaacacggagataaccaaagcgccacttagcgcctccatgatcaagaggtacgacgagcatcaccaggatctgacccttctgaaggctttggttcgccagcaactccccgagaagtacaaggagattttctttgaccaatcgaagaatggctacgcagggtacattgatggaggtaagttgttacttatgattgttttcctctctgctacatgtattttgttgttcatttctgtaagatataagaattgagttttcctctgatgatattattaggtgcaagtcaggaggaattctacaaattcatcaagcctattctggaaaagatggacggtacagaggagctgctcgttaaattgaaccgcgaagatttgcttcggaagcagcgtaccttcgacaatggcagcataccgcaccagatccacctcggtgagctgcatgctatcttgaggaggcaagaggacttctatccgttcctgaaagacaacagagagaagattgaaaagatcctcacgttccgcattccctactatgtaggttagtatcatatgaagaaatacctagtttcagttgatgaatgctattttctgacctcagttgttctcttttgagaattatttcttttctaatttgcctgatttttctattaattcattaggtccactcgcacgcgggaactcgcggtttgcgtggatgacacgcaaatccgaggagactatcacgccttggaacttcgaagaggtcgtggacaagggtgcgagtgcacagtccttcatcgaaaggatgaccaacttcgataagaatctcccaaatgagaaagtcctgcccaagcatagtctcctgtacgaatacttcacggtctacaacgagctgacgaaggtgaaatatgtgacggaggggatgcgcaaaccggccttcctgtcaggtaaatcctggtccacacttttacgataaaaacacaagattttaaactatgaactgatcaataatcattcctaaaagaccacacttttgttttgtttctaaagtaatttttactgttataacaggtgagcagaagaaggccattgtcgatctcttgttcaaaaccaatcggaaggtcactgtgaaacagcttaaagaggactactttaagaagatcgaatgctttgattctgtggaaatcagcggcgttgaggataggttcaatgcctctcttggcacataccatgacctgttgaaaatcatcaaggacaaggacttccttgacaacgaggagaacgaggacatcctcgaggacatcgtgctgactctcacgctgtttgaggacagagaaatgatcgaggagcgccttaagacttatgcgcatctgttcgatgacaaggtcatgaagcagttgaagaggaggagatatacaggtaagaggtcaaaaggtttccgcaatgatccctctttttttgtttctctagtttcaagaatttgggtatatgactaacttctgagtgttccttgatgcatatttgtgatgagacaaatgtttgttctatgttttaggttggggaaggctctccaggaagctcatcaacggcatccgcgacaagcaatccggcaagactatactggactttctcaaatccgacggttttgcgaatcggaacttcatgcagcttattcacgatgactcactgaccttcaaagaagatatccagaaggcccaagtgtcaggtcagggcgatagccttcacgaacacatagccaacctggctggatcgccagctataaagaagggcatactgcagacagtgaaggttgtggatgagctggtgaaggtaagttctgcatttggttatgctccttgcattttaggtgttcgtcgcacttccatttccatgaatagctaagattttttttctctgcattcattcttcttgcctcagttctaactgtttgtggtatttttgttttaattattgctacaggtcatgggccgccataagccggagaacatcgtcatcgagatggcgagggaaaaccagacgactcagaaagggcagaagaactcacgggagcgcatgaagcggatagaggaaggcatcaaggagcttgggagtcagattctgaaagagcacccagtcgaaaatactcaactccagaacgagaagctgtacctctattacctccagaatgggagagatatgtacgtcgaccaagagctcgacattaacagactctccgactatgatgtggatcacattgtccctcaatctttcctgaaggacgatagtattgacaacaaggtaaagcaactgtgttttaatcaatttcttgtcaggatatatggattataacttaatttttgagaaatctgtagtatttggcgtgaaatgagtttgctttttggtttctcccgtgttataggtccttacgcgctcagacaagaaccgcggaaaatccgacaatgtacccagcgaggaggttgtgaagaagatgaagaactattggaggcagcttttgaatgctaagctcataacccaacggaaattcgacaatctcacgaaggcagaaaggggcggactgtctgagctcgacaaagccggcttcatcaagcgccagttggttgaaactcgtcagattacgaaacatgtggcccagatactcgattcgcgtatgaatacgaagtatgatgagaatgacaaacttatcagggaggtaaaggtaaagtttccaactttcctttaccatatcaaactaaagttcgaaactttttatttgatcaacttcaaggccacccgatctttctattcctgattaatttgtgatgaatccatattgacttttgatggttacgcaggtgatcaccctcaagagcaaactggttagtgacttccggaaggacttccagttttacaaggttcgcgagatcaacaactaccatcatgcccatgacgcctacctgaacgccgttgttggcactgctctcatcaagaagtatccgaaactggagtctgagtttgtgtacggggattacaaggtgtacgacgttaggaagatgatcgcgaagtcagaacaagagatcggcaaggctaccgcgaaatacttcttttactcgaatatcatgaacttcttcaagacagagatcactctggcgaatggtgaaatccggaagaggcctctgatcgagacaaatggcgaaacaggtctgtctttcctatttcatatgtttaatcctaggaatttgatcaattgattgtatgtatgtcgatcccaagactttcttgttcacttatatcttaactctctctttgctgtttcttgcaggtgagattgtctgggataagggcagggattttgcgactgtgcgtaaggttctcagcatgccccaagtcaacatagtcaagaaaacggaggttcaaaccggtggtttctccaaggagtccattctccctaagcgcaactccgacaaactgattgcgaggaagaaggattgggatccgaagaaatacggaggctttgatagccctaccgtggcatacagcgtactggtagtggccaaggtggagaagggcaagagcaagaaactgaaaagcgtcaaggaactgcttggaattaccataatggaaaggtcctcgttcgagaagaatccgatcgacttcctcgaggctaaaggtaaaatattggatgccagacgatattctttcttttgatttgtaactttttcctgtcaaggtcgataaattttattttttttggtaaaaggtcgataatttttttttggagccattatgtaattttcctaattaactgaaccaaaattatacaaaccaggttacaaagaggtgaagaaagacctcattatcaaactgcccaagtattcgcttttcgaattggaaaatggcagaaaacgcatgctggcatctgccggagaactgcagaagggcaacgagctggcattgcccagtaagtacgtcaacttcctgtacttggcctcacactatgagaagctgaaggggtcaccagaggacaacgagcagaagcagttgtttgtcgagcagcacaagcactatcttgatgagatcatagagcagatcagcgaattttccaagcgggtcattcttgcagacgctaacctcgataaggtaaggacttctcatgaatattagtggcagattagtgttgttaaagtctttggttagataatcgatgcctcctaattgtccatgttttactggttttctacaattaaaggtgctttccgcgtacaacaagcacagagataagccgataagggaacaagcggaaaacatcatccacctgttcacactgaccaatctgggagccccagcagcctttaagtacttcgataccactatcgacagaaagcgctacacatcaaccaaggaagtgttggacgctacccttattcaccaatctattacagggctctatgagacaaggatagatctgtcgcagttgggtggtgactctagggctgacccaaagaagaagcgtaaagtctgagcttgctttcgttcgtatcatcggtttcgacaacgttcgtcaagttcaatgcatcagtttcattgcgcacacaccagaatcctactgagtttgagtattatggcattgggaaaactgtttttcttgtaccatttgttgtgcttgtaatttactgtgttttttattcggttttcgctatcgaactgtgaaatggaaatggatggagaagagttaatgaatgatatggtccttttgttcattctcaaattaatattatttgttttttctcttatttgttgtgtgttgaatttgaaattataagagatatgcaaacattttgttttgagtaaaaatgtgtcaaatcgtggcctctaatgaccgaagttaatatgaggagtaaaacacttgtagttgtaccattatgcttattcactaggcaacaaatatattttcagacctagaaaagctgcaaatgttactgaatacaagtatgtcctcttgtgttttagacatttatgaactttcctttatgtaattttccagaatccttgtcagattctaatcattgctttataattatagttatactcatggatttgtagttgagtatgaaaatattttttaatgcattttatgacttgccaattgattgacaac

**S5 tt*At*Cas12a cassette:**

*At*Ubi10 promoter

tt*At*Cas12a CDS

*Pisum sativum* E9 terminator

D156R

GGAGGTCGAGCTGCAGGTCAACGGATCAGGATATTCTTGTTTAAGATGTTGAACTCTATGGAGGTTTGTATGAACTGATGATCTAGGACCGGATAAGTTCCCTTCTTCATAGCGAACTTATTCAAAGAATGTTTTGTGTATCATTCTTGTTACATTGTTATTAATGAAAAAATATTATTGGTCATTGGACTGAACACGAGTGTTAAATATGGACCAGGCCCCAAATAAGATCCATTGATATATGAATTAAATAACAAGAATAAATCGAGTCACCAAACCACTTGCCTTTTTTAACGAGACTTGTTCACCAACTTGATACAAAAGTCATTATCCTATGCAAATCAATAATCATACAAAAATATCCAATAACACTAAAAAATTAAAAGAAATGGATAATTTCACAATATGTTATACGATAAAGAAGTTACTTTTCCAAGAAATTCACTGATTTTATAAGCCCACTTGCATTAGATAAATGGCAAAAAAAAACAAAAAGGAAAAGAAATAAAGCACGAAGAATTCTAGAAAATACGAAATACGCTTCAATGCAGTGGGACCCACGGTTCAATTATTGCCAATTTTCAGCTCCACCGTATATTTAAAAAATAAAACGATAATGCTAAAAAAATATAAATCGTAACGATCGTTAAATCTCAACGGCTGGATCTTATGACGACCGTTAGAAATTGTGGTTGTCGACGAGTCAGTAATAAACGGCGTCAAAGTGGTTGCAGCCGGCACACACGAGTCGTGTTTATCAACTCAAAGCACAAATACTTTTCCTCAACCTAAAAATAAGGCAATTAGCCAAAAACAACTTTGCGTGTAAACAACGCTCAATACACGTGTCATTTTATTATTAGCTATTGCTTCACCGCCTTAGCTTTCTCGTGACCTAGTCGTCCTCGTCTTTTCTTCTTCTTCTTCTATAAAACAATACCCAAAGAGCTCTTCTTCTTCACAATTCAGATTTCAATTTCTCAAAATCTTAAAAACTTTCTCTCAATTCTCTCTACCGTGATCAAGGTAAATTTCTGTGTTCCTTATTCTCTCAAAATCTTCGATTTTGTTTTCGTTCGATCCCAATTTCGTATATGTTCTTTGGTTTAGATTCTGTTAATCTTAGATCGAACACGATTTTCTGGGTTTGATCGTTAGATATCATCTTAATTCTCGATTAGGGTTTCATAGATATCATCCGATTTGTTCAAATAATTTGAGTTTTGTCGAATAATTACTCTTCGATTTGTGATTTCTATCTAGATCTGGTGTTAGTTTCTAGTTTGTGCGATCGAATTTGTCGATTAATCTGAGTTTTTCTGATTAACAGGaatgagcaagctcgagaagtttaccaactgctacagcctctctaagaccctcaggttcaaggctatccctgtgggaaagacccaagagaatatcgacaacaagaggctcctcgtcgaggatgagaagagagctgaagattacaagggcgtgaagaagctcctcgacaggtactacctcagcttcatcaacgatgtgctccacagcatcaagctcaagaacctcaacaactacatcagcctcttccgtaagaaaaccaggaccgagaaagagaacaaagagcttgagaacctcgagatcaacctccgtaaagagatcgccaaggctttcaagggaaacgagggatacaagagcctcttcaagaaggatattatcgagacaatcctgcctgagttcctggacgataaggatgagatcgctctcgtgaacagcttcaacggattcactactgccttcaccggattcttcagaaacagggaaaacatgttcagcgaagaggccaagagcacctctatcgctttcagatgcatcaacgagaacctcacgcgttacatcagcaacatggacatcttcgagaaggtggacgccatcttcgataagcacgaggtgcaagaaatcaaagagaagatcctcaacagcgactacgacgtcgaggacttttttgaaggggagttcttcaacttcgttctcacccaagagggcatcgacgtgtacaacgctattatcggaggattcgtgaccgagtctggggagaagattaagggactcaacgagtacatcaacctgtacaaccagaaaacgaagcagaagctcccgaagttcaagccgctctacaagcaggttctctctgatcgtgagagcctctcattttacggtgagggttacacctctgacgaggaagtgcttgaggttttccgtaacaccctcaacaagaacagcgagatcttctcgtccatcaagaagttggagaagcttttcaagaacttcgacgagtacagcagcgctgggatcttcgttaagaacggacctgctatcagcaccatcagcaaggatattttcggcgagtggaacgtgatcagggacaagtggaatgctgagtacgatgacatccacctcaagaagaaggctgtcgtcactgagaagtacgaggatgacaggcgtaagtcgttcaagaagatcggctctttcagcctcgagcagcttcaagaatacgctgatgctgatctcagcgtggtcgagaagctcaaagagatcatcatccagaaggtcgacgagatctacaaggtgtacgggtcctctgagaagttgttcgatgctgatttcgtcctcgagaagagtctgaagaagaacgacgctgtcgtcgcgatcatgaaggatttgctcgacagcgtgaagtccttcgagaactatatcaaggccttcttcggagagggcaaagagactaatagggacgagtctttctacggggatttcgtgctcgcttacgatatcctcctcaaggtggaccatatctacgacgccatcagaaactacgtgacccagaagccttacagcaaggacaagttcaagttgtactttcagaacccgcagttcatgggcggatgggacaaagacaaagagacagattacagggccaccatcctcaggtacgggtctaagtactacctggccatcatggacaagaaatacgccaagtgcctccaaaagatcgacaaggatgacgtgaacgggaactatgagaagatcaactacaagctccttccgggaccgaacaagatgcttcctaaggtgttcttcagcaagaaatggatggcctactacaacccgtctgaggacatccagaaaatctacaagaacgggaccttcaagaaaggcgacatgttcaacctcaacgactgccacaagctcatcgatttcttcaaggacagcatctcgcgttacccgaagtggtctaacgcttacgactttaacttcagcgagacagaaaagtacaaggatatcgccgggttctaccgtgaggttgaggaacagggttacaaggttagcttcgagagcgcctccaagaaagaggttgacaagttggtcgaagagggcaagctctacatgttccagatctataacaaggacttctccgacaagagccacggaactcctaacctccatacgatgtacttcaagctgcttttcgacgagaacaaccacgggcagatcagactttctggtggtgctgaactcttcatgcgtagggcctcactcaagaaagaagagttggttgttcacccggccaactctccaatcgctaacaagaatcctgacaacccgaaaaagaccaccacgctgtcttacgacgtctacaaggacaaaaggttcagcgaggaccagtacgagcttcatatcccgatcgctatcaacaagtgcccgaagaacatcttcaagatcaataccgaggtgagggtgctgctcaagcacgatgataacccttacgtgatcggaatcgatcgtggtgagagaaacctcctctacatcgttgtggtggacggaaagggaaacatcgtcgagcagtacagcctgaacgagattatcaacaatttcaacggcatcaggatcaagaccgactaccactcactcctcgataagaaagaaaaagagcgtttcgaggccaggcagaactggacttctatcgaaaacatcaaagagttgaaggccggctacatctctcaggtggtgcataagatctgcgagctggtggaaaagtacgatgctgtgatcgctcttgaggacctcaactctgggttcaagaacagtagagtgaaggttgagaagcaggtctaccaaaagttcgagaagatgctcatcgacaagctcaactacatggtggacaaaaagagcaacccttgcgctaccggtggtgctcttaagggataccagatcacgaacaagttcgagtccttcaagagcatgagcacccagaacggcttcatcttctatatccctgcttggctcaccagcaagatcgatccttctactggtttcgtgaacctgctcaagaccaagtacacctcgatcgccgacagcaagaagttcatctcgtctttcgacaggatcatgtacgtgccggaagaggatcttttcgagttcgctctcgactataagaacttcagcaggaccgacgccgactacattaagaagtggaagctctactcctacgggaaccgtatcaggatcttccgaaatccgaagaaaaacaacgtgttcgactgggaagaagtgtgcctcacctctgcctacaaagaactgttcaacaagtacggcatcaactaccagcagggtgatatcagggctcttttgtgcgagcagagcgacaaggcattctacagctcattcatggccctcatgtctctcatgctccagatgaggaactctatcaccggaaggaccgatgtggacttccttatctctccggtcaagaactctgacgggatcttctacgacagccgtaactatgaggctcaagagaacgctatcctgccgaagaatgctgatgcaaacggggcttacaacattgcgagaaaggttctctgggctatcgggcagtttaagaaagcggaagatgagaagctggacaaggtgaagatcgccatctccaacaaagagtggcttgagtacgctcagacctccgttaagcacaagaggcctgctgctactaagaaagctggtcaggctaagaagaagaaatgagcttgctttcgttcgtatcatcggtttcgacaacgttcgtcaagttcaatgcatcagtttcattgcgcacacaccagaatcctactgagtttgagtattatggcattgggaaaactgtttttcttgtaccatttgttgtgcttgtaatttactgtgttttttattcggttttcgctatcgaactgtgaaatggaaatggatggagaagagttaatgaatgatatggtccttttgttcattctcaaattaatattatttgttttttctcttatttgttgtgtgttgaatttgaaattataagagatatgcaaacattttgttttgagtaaaaatgtgtcaaatcgtggcctctaatgaccgaagttaatatgaggagtaaaacacttgtagttgtaccattatgcttattcactaggcaacaaatatattttcagacctagaaaagctgcaaatgttactgaatacaagtatgtcctcttgtgttttagacatttatgaactttcctttatgtaattttccagaatccttgtcagattctaatcattgctttataattatagttatactcatggatttgtagttgagtatgaaaatattttttaatgcattttatgacttgccaattgattgacaac

**S5 V1 guide cassette:**

*At*U626 promoter

*Lb*Cas12a DR invariable guide sequence

Variable spacer sequence

HDV ribozyme

Terminator

catcttcattcttaagatatgaagataatcttcaaaaggcccctgggaatctgaaagaagagaagcaggcccatttatatgggaaagaacaatagtatttcttatataggcccatttaagttgaaaacaatcttcaaaagtcccacatcgcttagataagaaaacgaagctgagtttatatacagctagagtcgaagtagtgattgtaatttctactaagtgtagatNNNNNNNNNNNNNNNNNNNNNNNtaatttctactaagtgtagatNNNNNNNNNNNNNNNNNNNNNNNtaatttctactaagtgtagatNNNNNNNNNNNNNNNNNNNNNNNtaatttctactaagtgtagatNNNNNNNNNNNNNNNNNNNNNNNtaatttctactaagtgtagatgtcccttcgaagggcaattctgcagatatccatcacactggcggccgctcgaggtcgagggtatcgataagcttttttttttt

**S6 guide cassette:**

*At*U626 promoter

HH ribozyme

*Lb*Cas12a DR invariable guide sequence

Variable spacer sequence

HDV ribozyme

Terminator

CATCTTCATTCTTAAGATATGAAGATAATCTTCAAAAGGCCCCTGGGAATCTGAAAGAAGAGAAGCAGGCCCATTTATATGGGAAAGAACAATAGTATTTCTTATATAGGCCCATTTAAGTTGAAAACAATCTTCAAAAGTCCCACATCGCTTAGATAAGAAAACGAAGCTGAGTTTATATACAGCTAGAGTCGAAGTAGTGATTGaaattactgatgagtccgtgaggacgaaacgagtaagctcgtctaatttctactaagtgtagatNNNNNNNNNNNNNNNNNNNNNNNggccggcatggtcccagcctcctcgctggcgccggctgggcaacatgcttcggcatggcgaatgggacttttt

**S7 tt*Hs*Cas12a cassette:**

*At*Ubi10 promoter

tt*Hs*Cas12a CDS

*Pisum sativum* E9 terminator

D156R

GTCGAGCTGCAGGTCAACGGATCAGGATATTCTTGTTTAAGATGTTGAACTCTATGGAGGTTTGTATGAACTGATGATCTAGGACCGGATAAGTTCCCTTCTTCATAGCGAACTTATTCAAAGAATGTTTTGTGTATCATTCTTGTTACATTGTTATTAATGAAAAAATATTATTGGTCATTGGACTGAACACGAGTGTTAAATATGGACCAGGCCCCAAATAAGATCCATTGATATATGAATTAAATAACAAGAATAAATCGAGTCACCAAACCACTTGCCTTTTTTAACGAGACTTGTTCACCAACTTGATACAAAAGTCATTATCCTATGCAAATCAATAATCATACAAAAATATCCAATAACACTAAAAAATTAAAAGAAATGGATAATTTCACAATATGTTATACGATAAAGAAGTTACTTTTCCAAGAAATTCACTGATTTTATAAGCCCACTTGCATTAGATAAATGGCAAAAAAAAACAAAAAGGAAAAGAAATAAAGCACGAAGAATTCTAGAAAATACGAAATACGCTTCAATGCAGTGGGACCCACGGTTCAATTATTGCCAATTTTCAGCTCCACCGTATATTTAAAAAATAAAACGATAATGCTAAAAAAATATAAATCGTAACGATCGTTAAATCTCAACGGCTGGATCTTATGACGACCGTTAGAAATTGTGGTTGTCGACGAGTCAGTAATAAACGGCGTCAAAGTGGTTGCAGCCGGCACACACGAGTCGTGTTTATCAACTCAAAGCACAAATACTTTTCCTCAACCTAAAAATAAGGCAATTAGCCAAAAACAACTTTGCGTGTAAACAACGCTCAATACACGTGTCATTTTATTATTAGCTATTGCTTCACCGCCTTAGCTTTCTCGTGACCTAGTCGTCCTCGTCTTTTCTTCTTCTTCTTCTATAAAACAATACCCAAAGAGCTCTTCTTCTTCACAATTCAGATTTCAATTTCTCAAAATCTTAAAAACTTTCTCTCAATTCTCTCTACCGTGATCAAGGTAAATTTCTGTGTTCCTTATTCTCTCAAAATCTTCGATTTTGTTTTCGTTCGATCCCAATTTCGTATATGTTCTTTGGTTTAGATTCTGTTAATCTTAGATCGAACACGATTTTCTGGGTTTGATCGTTAGATATCATCTTAATTCTCGATTAGGGTTTCATAGATATCATCCGATTTGTTCAAATAATTTGAGTTTTGTCGAATAATTACTCTTCGATTTGTGATTTCTATCTAGATCTGGTGTTAGTTTCTAGTTTGTGCGATCGAATTTGTCGATTAATCTGAGTTTTTCTGATTAACAGGAATGagcaagctggagaagtttacaaactgctactccctgtctaagaccctgaggttcaaggccatccctgtgggcaagacccaggagaacatcgacaataagcggctgctggtggaggacgagaagagagccgaggattataagggcgtgaagaagctgctggatcgctactatctgtcttttatcaacgacgtgctgcacagcatcaagctgaagaatctgaacaattacatcagcctgttccggaagaaaaccagaaccgagaaggagaataaggagctggagaacctggagatcaatctgcggaaggagatcgccaaggccttcaagggcaacgagggctacaagtccctgtttaagaaggatatcatcgagacaatcctgccagagttcctggacgataaggacgagatcgccctggtgaacagcttcaatggctttaccacagccttcaccggcttctttCGCaacagagagaatatgttttccgaggaggccaagagcacatccatcgccttcaggtgtatcaacgagaatctgacccgctacatctctaatatggacatcttcgagaaggtggacgccatctttgataagcacgaggtgcaggagatcaaggagaagatcctgaacagcgactatgatgtggaggatttctttgagggcgagttctttaactttgtgctgacacaggagggcatcgacgtgtataacgccatcatcggcggcttcgtgaccgagagcggcgagaagatcaagggcctgaacgagtacatcaacctgtataatcagaaaaccaagcagaagctgcctaagtttaagccactgtataagcaggtgctgagcgatcgggagtctctgagcttctacggcgagggctatacatccgatgaggaggtgctggaggtgtttagaaacaccctgaacaagaacagcgagatcttcagctccatcaagaagctggagaagctgttcaagaattttgacgagtactctagcgccggcatctttgtgaagaacggccccgccatcagcacaatctccaaggatatcttcggcgagtggaacgtgatccgggacaagtggaatgccgagtatgacgatatccacctgaagaagaaggccgtggtgaccgagaagtacgaggacgatcggagaaagtccttcaagaagatcggctccttttctctggagcagctgcaggagtacgccgacgccgatctgtctgtggtggagaagctgaaggagatcatcatccagaaggtggatgagatctacaaggtgtatggctcctctgagaagctgttcgacgccgattttgtgctggagaagagcctgaagaagaacgacgccgtggtggccatcatgaaggacctgctggattctgtgaagagcttcgagaattacatcaaggccttctttggcgagggcaaggagacaaacagggacgagtccttctatggcgattttgtgctggcctacgacatcctgctgaaggtggaccacatctacgatgccatccgcaattatgtgacccagaagccctactctaaggataagttcaagctgtattttcagaaccctcagttcatgggcggctgggacaaggataaggagacagactatcgggccaccatcctgagatacggctccaagtactatctggccatcatggataagaagtacgccaagtgcctgcagaagatcgacaaggacgatgtgaacggcaattacgagaagatcaactataagctgctgcccggccctaataagatgctgccaaaggtgttcttttctaagaagtggatggcctactataaccccagcgaggacatccagaagatctacaagaatggcacattcaagaagggcgatatgtttaacctgaatgactgtcacaagctgatcgacttctttaaggatagcatctcccggtatccaaagtggtccaatgcctacgatttcaacttttctgagacagagaagtataaggacatcgccggcttttacagagaggtggaggagcagggctataaggtgagcttcgagtctgccagcaagaaggaggtggataagctggtggaggagggcaagctgtatatgttccagatctataacaaggacttttccgataagtctcacggcacacccaatctgcacaccatgtacttcaagctgctgtttgacgagaacaatcacggacagatcaggctgagcggaggagcagagctgttcatgaggcgcgcctccctgaagaaggaggagctggtggtgcacccagccaactcccctatcgccaacaagaatccagataatcccaagaaaaccacaaccctgtcctacgacgtgtataaggataagaggttttctgaggaccagtacgagctgcacatcccaatcgccatcaataagtgccccaagaacatcttcaagatcaatacagaggtgcgcgtgctgctgaagcacgacgataacccctatgtgatcggcatcgataggggcgagcgcaatctgctgtatatcgtggtggtggacggcaagggcaacatcgtggagcagtattccctgaacgagatcatcaacaacttcaacggcatcaggatcaagacagattaccactctctgctggacaagaaggagaaggagaggttcgaggcccgccagaactggacctccatcgagaatatcaaggagctgaaggccggctatatctctcaggtggtgcacaagatctgcgagctggtggagaagtacgatgccgtgatcgccctggaggacctgaactctggctttaagaatagccgcgtgaaggtggagaagcaggtgtatcagaagttcgagaagatgctgatcgataagctgaactacatggtggacaagaagtctaatccttgtgcaacaggcggcgccctgaagggctatcagatcaccaataagttcgagagctttaagtccatgtctacccagaacggcttcatcttttacatccctgcctggctgacatccaagatcgatccatctaccggctttgtgaacctgctgaaaaccaagtataccagcatcgccgattccaagaagttcatcagctcctttgacaggatcatgtacgtgcccgaggaggatctgttcgagtttgccctggactataagaacttctctcgcacagacgccgattacatcaagaagtggaagctgtactcctacggcaaccggatcagaatcttccggaatcctaagaagaacaacgtgttcgactgggaggaggtgtgcctgaccagcgcctataaggagctgttcaacaagtacggcatcaattatcagcagggcgatatcagagccctgctgtgcgagcagtccgacaaggccttctactctagctttatggccctgatgagcctgatgctgcagatgcggaacagcatcacaggccgcaccgacgtggattttctgatcagccctgtgaagaactccgacggcatcttctacgatagccggaactatgaggcccaggagaatgccatcctgccaaagaacgccgacgccaatggcgcctataacatcgccagaaaggtgctgtgggccatcggccagttcaagaaggccgaggacgagaagctggataaggtgaagatcgccatctctaacaaggagtggctggagtacgcccagaccagcgtgaagcacaaaaggccggcggccacgaaaaaggccggccaggcaaaaaagaaaaagtaggcttgctttcgttcgtatcatcggtttcgacaacgttcgtcaagttcaatgcatcagtttcattgcgcacacaccagaatcctactgagtttgagtattatggcattgggaaaactgtttttcttgtaccatttgttgtgcttgtaatttactgtgttttttattcggttttcgctatcgaactgtgaaatggaaatggatggagaagagttaatgaatgatatggtccttttgttcattctcaaattaatattatttgttttttctcttatttgttgtgtgttgaatttgaaattataagagatatgcaaacattttgttttgagtaaaaatgtgtcaaatcgtggcctctaatgaccgaagttaatatgaggagtaaaacacttgtagttgtaccattatgcttattcactaggcaacaaatatattttcagacctagaaaagctgcaaatgttactgaatacaagtatgtcctcttgtgttttagacatttatgaactttcctttatgtaattttccagaatccttgtcagattctaatcattgctttataattatagttatactcatggatttgtagttgagtatgaaaatattttttaatgcattttatgacttgccaattgattgacaac

**S8 tt*At*Cas12a+int with 8 introns cassette:**

tt*At*Ubi10 promoter

*At*Cas12a CDS with 8 introns

*Pisum sativum* E9 terminator

Intron

D156R

GGAGGTCGAGCTGCAGGTCAACGGATCAGGATATTCTTGTTTAAGATGTTGAACTCTATGGAGGTTTGTATGAACTGATGATCTAGGACCGGATAAGTTCCCTTCTTCATAGCGAACTTATTCAAAGAATGTTTTGTGTATCATTCTTGTTACATTGTTATTAATGAAAAAATATTATTGGTCATTGGACTGAACACGAGTGTTAAATATGGACCAGGCCCCAAATAAGATCCATTGATATATGAATTAAATAACAAGAATAAATCGAGTCACCAAACCACTTGCCTTTTTTAACGAGACTTGTTCACCAACTTGATACAAAAGTCATTATCCTATGCAAATCAATAATCATACAAAAATATCCAATAACACTAAAAAATTAAAAGAAATGGATAATTTCACAATATGTTATACGATAAAGAAGTTACTTTTCCAAGAAATTCACTGATTTTATAAGCCCACTTGCATTAGATAAATGGCAAAAAAAAACAAAAAGGAAAAGAAATAAAGCACGAAGAATTCTAGAAAATACGAAATACGCTTCAATGCAGTGGGACCCACGGTTCAATTATTGCCAATTTTCAGCTCCACCGTATATTTAAAAAATAAAACGATAATGCTAAAAAAATATAAATCGTAACGATCGTTAAATCTCAACGGCTGGATCTTATGACGACCGTTAGAAATTGTGGTTGTCGACGAGTCAGTAATAAACGGCGTCAAAGTGGTTGCAGCCGGCACACACGAGTCGTGTTTATCAACTCAAAGCACAAATACTTTTCCTCAACCTAAAAATAAGGCAATTAGCCAAAAACAACTTTGCGTGTAAACAACGCTCAATACACGTGTCATTTTATTATTAGCTATTGCTTCACCGCCTTAGCTTTCTCGTGACCTAGTCGTCCTCGTCTTTTCTTCTTCTTCTTCTATAAAACAATACCCAAAGAGCTCTTCTTCTTCACAATTCAGATTTCAATTTCTCAAAATCTTAAAAACTTTCTCTCAATTCTCTCTACCGTGATCAAGGTAAATTTCTGTGTTCCTTATTCTCTCAAAATCTTCGATTTTGTTTTCGTTCGATCCCAATTTCGTATATGTTCTTTGGTTTAGATTCTGTTAATCTTAGATCGAACACGATTTTCTGGGTTTGATCGTTAGATATCATCTTAATTCTCGATTAGGGTTTCATAGATATCATCCGATTTGTTCAAATAATTTGAGTTTTGTCGAATAATTACTCTTCGATTTGTGATTTCTATCTAGATCTGGTGTTAGTTTCTAGTTTGTGCGATCGAATTTGTCGATTAATCTGAGTTTTTCTGATTAACAGGaatgagcaagctcgagaagtttaccaactgctacagcctctctaagaccctcaggttcaaggctatccctgtgggaaagacccaagagaatatcgacaacaagaggctcctcgtcgaggatgagaagagagctgaagattacaagggcgtgaagaagctcctcgacaggtactacctcagcttcatcaacgatgtgctccacagcatcaagctcaagaacctcaacaactacatcagcctcttccgtaagaaaaccaggaccgagaaagagaacaaagagcttgagaacctcgagatcaacctccgtaaagagatcgccaaggctttcaagggaaacgagggatacaagagcctcttcaagaaggatattatcgagacaatcctgcctgagttcctggacgataaggatgagatcgctctcgtgaacagcttcaacggattcactactgccttcaccggattcttcagaaacagggaaaacatgttcagcgaagaggccaagagcacctctatcgctttcagatgcatcaacgagaacctcacgcgttacatcagcaacatggacatcttcgagaaggtaacattccttagttacctttcttttctttttccatcataagtttatagattgtacatgctttgagatttttctttgcaaacaatctcaggtggacgccatcttcgataagcacgaggtgcaagaaatcaaagagaagatcctcaacagcgactacgacgtcgaggacttttttgaaggggagttcttcaacttcgttctcacccaagagggcatcgacgtgtacaacgctattatcggaggattcgtgaccgagtctggggagaagattaagggactcaacgagtacatcaacctgtacaaccagaaaacgaagcagaagctcccgaagttcaagccgctctacaagcaggtctgtctttcctatttcatatgtttaatcctaggaatttgatcaattgattgtatgtatgtcgatcccaagactttcttgttcacttatatcttaactctctctttgctgtttcttgcaggttctctctgatcgtgagagcctctcattttacggtgagggttacacctctgacgaggaagtgcttgaggttttccgtaacaccctcaacaagaacagcgagatcttctcgtccatcaagaagttggagaagcttttcaagaacttcgacgagtacagcagcgctgggatcttcgttaagaacggacctgctatcagcaccatcagcaaggatattttcggcgagtggaacgtgatcagggacaagtggaatgctgagtacgatgacatccacctcaagaagaaggctgtcgtcactgagaagtacgaggatgacaggcgtaagtcgttcaagaagatcggctctttcagcctcgagcagcttcaagaatacgctgatgctgatctcagcgtggtcgagaagctcaaagagatcatcatccagaaggtcgacgagatctacaaggtaagttgttacttatgattgttttcctctctgctacatgtattttgttgttcatttctgtaagatataagaattgagttttcctctgatgatattattaggtgtacgggtcctctgagaagttgttcgatgctgatttcgtcctcgagaagagtctgaagaagaacgacgctgtcgtcgcgatcatgaaggatttgctcgacagcgtgaagtccttcgagaactatatcaaggccttcttcggagagggcaaagagactaatagggacgagtctttctacggggatttcgtgctcgcttacgatatcctcctcaaggtggaccatatctacgacgccatcagaaactacgtgacccagaagccttacagcaaggacaagttcaagttgtactttcagaacccgcagttcatgggcggatgggacaaagacaaagagacagattacagggccaccatcctcaggttagtatcatatgaagaaatacctagtttcagttgatgaatgctattttctgacctcagttgttctcttttgagaattatttcttttctaatttgcctgatttttctattaattcattaggtacgggtctaagtactacctggccatcatggacaagaaatacgccaagtgcctccaaaagatcgacaaggatgacgtgaacgggaactatgagaagatcaactacaagctccttccgggaccgaacaagatgcttcctaaggtgttcttcagcaagaaatggatggcctactacaacccgtctgaggacatccagaaaatctacaagaacgggaccttcaagaaaggcgacatgttcaacctcaacgactgccacaagctcatcgatttcttcaaggacagcatctcgcgttacccgaagtggtctaacgcttacgactttaacttcagcgagacagaaaagtacaaggatatcgccgggttctaccgtgaggttgaggaacagggttacaaAgttagcttcgagagcgcctccaagaaagaggtaaatcctggtccacacttttacgataaaaacacaagattttaaactatgaactgatcaataatcattcctaaaagaccacacttttgttttgtttctaaagtaatttttactgttataacaggttgacaagttggtcgaagagggcaagctctacatgttccagatctataacaaggacttctccgacaagagccacggaactcctaacctccatacgatgtacttcaagctgcttttcgacgagaacaaccacgggcagatcagactttctggtggtgctgaactcttcatgcgtagggcctcactcaagaaagaagagttggttgttcacccggccaactctccaatcgctaacaagaatcctgacaacccgaaaaagaccaccacgctgtcttacgacgtctacaaggacaaaaggttcagcgaggaccagtacgagcttcatatcccgatcgctatcaacaagtgcccgaagaacatcttcaagatcaataccgaggtaaggacttctcatgaatattagtggcagattagtgttgttaaagtctttggttagataatcgatgcctcctaattgtccatgttttactggttttctacaattaaaggtgagggtgctgctcaagcacgatgataacccttacgtgatcggaatcgatcgtggtgagagaaacctcctctacatcgttgtggtggacggaaagggaaacatcgtcgagcagtacagcctgaacgagattatcaacaatttcaacggcatcaggatcaagaccgactaccactcactcctcgataagaaagaaaaagagcgtttcgaggccaggcagaactggacttctatcgaaaacatcaaagagttgaaggccggctacatctctcaggtggtgcataagatctgcgagctggtggaaaagtacgatgctgtgatcgctcttgaggacctcaactctgggttcaagaacagtagagtgaaggtaagttctgcatttggttatgctccttgcattttaggtgttcgtcgcacttccatttccatgaatagctaagattttttttctctgcattcattcttcttgcctcagttctaactgtttgtggtatttttgttttaattattgctacaggttgagaagcaggtctaccaaaagttcgagaagatgctcatcgacaagctcaactacatggtggacaaaaagagcaacccttgcgctaccggtggtgctcttaagggataccagatcacgaacaagttcgagtccttcaagagcatgagcacccagaacggcttcatcttctatatccctgcttggctcaccagcaagatcgatccttctactggtttcgtgaacctgctcaagaccaagtacacctcgatcgccgacagcaagaagttcatctcgtctttcgacaggatcatgtacgtgccggaagaggatcttttcgagttcgctctcgactataagaacttcagcaggaccgacgccgactacattaagaagtggaagctctactcctacgggaaccgtatcaggatcttccgaaatccgaagaaaaacaacgtgttcgactgggaagaagtgtgcctcacctctgcctacaaagaactgttcaacaagtacggcatcaactaccagcagggtgatatcagggctcttttgtgcgagcagagcgacaaggcattctacagctcattcatggccctcatgtctctcatgctccagatgaggaactctatcaccggaaggaccgatgtggacttccttatctctccggtcaagaactctgacgggatcttctacgacagccgtaactatgaggctcaagagaacgctatcctgccgaagaatgctgatgcaaacggggcttacaacattgcgagaaaggtaaagcaactgtgttttaatcaatttcttgtcaggatatatggattataacttaatttttgagaaatctgtagtatttggcgtgaaatgagtttgctttttggtttctcccgtgttataggttctctgggctatcgggcagtttaagaaagcggaagatgagaagctggacaaggtgaagatcgccatctccaacaaagagtggcttgagtacgctcagacctccgttaagcacaagaggcctgctgctactaagaaagctggtcaggctaagaagaagaaatgagcttgctttcgttcgtatcatcggtttcgacaacgttcgtcaagttcaatgcatcagtttcattgcgcacacaccagaatcctactgagtttgagtattatggcattgggaaaactgtttttcttgtaccatttgttgtgcttgtaatttactgtgttttttattcggttttcgctatcgaactgtgaaatggaaatggatggagaagagttaatgaatgatatggtccttttgttcattctcaaattaatattatttgttttttctcttatttgttgtgtgttgaatttgaaattataagagatatgcaaacattttgttttgagtaaaaatgtgtcaaatcgtggcctctaatgaccgaagttaatatgaggagtaaaacacttgtagttgtaccattatgcttattcactaggcaacaaatatattttcagacctagaaaagctgcaaatgttactgaatacaagtatgtcctcttgtgttttagacatttatgaactttcctttatgtaattttccagaatccttgtcagattctaatcattgctttataattatagttatactcatggatttgtagttgagtatgaaaatattttttaatgcattttatgacttgccaattgattgacaac
